# Age and dose-dependent effects of alpha-lipoic acid on human microtubule-associated protein tau-induced endoplasmic reticulum unfolded protein response: implications for Alzheimer’s disease

**DOI:** 10.1101/2020.07.31.230847

**Authors:** Elahe Zarini-Gakiye, Nima Sanadgol, Kazem Parivar, Gholamhassan Vaezi

**Affiliations:** Department of Biology, Science and Research Branch, Islamic Azad University, Tehran, Iran; Department of Biology, Faculty of Sciences, University of Zabol, Zabol, Iran; Department of Biology, Damghan Branch, Islamic Azad University, Damghan, Semnan, Iran

**Keywords:** Aging, Alzheimer’s disease, IRE1-XBP1 pathway, PERK-ATF4 pathway

## Abstract

**Background:** In human tauopathies, pathological aggregation of misfolded/unfolded proteins particularly microtubule-associated protein tau (MAPT, tau) is considered to be essential mechanisms that trigger the induction of endoplasmic reticulum (ER) stress. Here we assessed the molecular effects of natural antioxidant alpha-lipoic acid (ALA) in human tau^R406W^ (htau)-induced ER unfolded protein response (ERUPR) in the young and older flies.

**Methods:** In order to reduce htau neurotoxicity during brain development, we used a transgenic model of tauopathy where the maximum toxicity was observed in adult flies. Then, the effects of ALA (0.001, 0.005, and 0.025% w/w of diet) in htau-induced ERUPR in the ages 20 and 30 days were evaluated.

**Results:** Data from expression (mRNA and protein) patterns of htau, analysis of eyes external morphology as well as larvae olfactory memory were confirmed our tauopathy model. Moreover, expression of ERUPR-related proteins involving activating transcription factor 6 (ATF6), inositol regulating enzyme 1 (IRE1), and protein kinase RNA-like ER kinase (PERK) were upregulated and locomotor function decreased in both ages of the model flies. Remarkably, the lower dose of ALA modified ERUPR and supported the reduction of behavioral deficits in youngest adults through enhancement of GRP87/Bip, reduction of ATF6, downregulation of PERK-ATF4 pathway, and activation of the IRE1-XBP1 pathway. On the other hand, only a higher dose of ALA was able to affect the ERUPR via moderation of PERK-ATF4 signaling in the oldest adults. As ALA exerts their higher protective effects on the locomotor function of younger adults when htau^R406W^ expressed in all neurons (htau-*elav*) and mushroom body neurons (htau-*ok*), we proposed that ALA has age-dependent effects in this model.

**Conclusion:** Taken together, based on our results we conclude that aging potentially influences the ALA effective dose and mechanism of action on tau-induced ERUPR. Further molecular studies will warrant possible therapeutic applications of ALA in age-related tauopathies.

## Background

Alzheimer’s disease (AD) is the main form of dementia among the elderly and the most prevalent neurodegenerative disorder. It represents the most important global health problem, particularly in developing and developed countries, due to the immense influence on persons, families, the health care system, and society in general [1]. Currently, over 47 million people are living with dementia worldwide and this amount is expected to reach 131 million in 2050 [2, 3]. People 65 years and older are more at risk as a target group, which puts aging as the important risk factors for disease progression [4, 5].

The classical neuropathology of AD involves the progressive accumulation of extracellular amyloid beta-protein (Aβ) plaques, derived by sequential proteolytic processing from the larger transmembrane amyloid precursor protein (APP), and intracellular filamentous neurofibrillary tangles (NFTs), which result from the aggregation of the soluble hyper-phosphorylated human microtubule-associated protein tau (hMAPT or htau) [6, 7]. In the normal situation, tau is a soluble and unstructured protein that is bound to microtubule to regulate its stability in the axon and this binding itself regulated by phosphorylation of its microtubule-binding domains. A high level of phosphorylated tau results in detachment of tau from the microtubule, its aggregation, and eventually the formation of NFTs in the neocortical and hippocampal areas [8]. It has confirmed that mutation in the htau gene including htau^R406W^, htau^V337M^, and hTau^P301L^ results in autosomal dominant neurodegenerative disease. These forms of mutations lead to hyper-phosphorylation and tau aggregation which is one of the well-known genetic factors underlying the progress of AD [9, 10]. Therefore, the determination of molecular pathways that change tau neurotoxicity could help to a better understanding of the pathogenesis of all tauopathies, including AD.

Frequently, NFTs induce endoplasmic reticulum (ER) stress and finally trigger the unfolded protein response (UPR) that may protect cells against the buildup of these toxic proteins in the neurons [11]. The ER microenvironment adversities like various forms of nutrient deprivation, changes in calcium balance, redox potential, profound hypoxia, and accumulation of misfolded/unfolded proteins potentially prompt the ERUPR. The ERUPR signaling pathway is highly conserved in all eukaryotic cells involved in the cell-homeostasis through the transcriptional remodeling of ER proteostasis□related pathways. The ERUPR is induced by three stress-responsive transmembrane proteins including activating transcription factor 6 (ATF6), inositol regulating enzyme 1 (IRE1), and protein kinase RNA-like ER kinase (PERK) [12, 13]. It has been demonstrated that there is a failure in the cell’s ability to handle folding, accumulation, and aggregation of newly synthesized proteins due to dysfunction of the UPR system during aging [14]. Therefore, reconstruction and recovery of the UPR proper function attract significant interest in elderly and aging-related neurodegenerative diseases.

Alpha-Lipoic acid [ALA, 5-(1,2-dithiolan-3-yl)-pentanoic acid] is a naturally occurring disulfide compound which advised as a supplement, possessing various neuroprotective, pharmacologic, and antioxidant activities [15, 16]. ALA synthesized enzymatically in mitochondria of humans, animals, and plants (in small amounts in spinach, broccoli, peas, Brussels sprouts, and potatoes) from octanoic acid and cysteine. Moreover, ALA is easily absorbed in the diet, transported, taken up by cells, and changed to the dihydrolipoic acid, the reduced form of ALA, in several tissues, including the brain [17]. It was showed that ALA induced ER-stress mediated DNA fragmentation and apoptosis in A549 cell lines in a time- and dose-dependent manner [18]. On the other hand, ALA prevented ER stress-induced insulin resistance in HepG2 cells by improving mitochondrial function [19]. ALA has also been shown to possess a range of benefits in other neurodegenerative diseases which are similar to principles of AD pathogenesis [20, 21]. Thus, ALA could be considered in designing new multi-target ligands as an excellent structure for the management, and possibly treatment of AD.

While senescence has been suggested as a potential risk factor for neurodegeneration, the biological significance of its signaling cascade on tauopathy-induced ERUPR and behavioral dysfunctions has not been comprehensively evaluated [22]. *Drosophila melanogaster* has emerged as a powerful tractable organism in which to model aspects of both human aging and tauopathies [23]. For simultaneous connecting between tauopathy and aging, we minimized htau^R406W^ toxicity during larval development and younger adults and maximized its toxicity in older adults (20 and 30 days old) via regulation of htau transcription in *Drosophila* neurons. With this adult-onset model, we reduced htau^R406W^-induced neurotoxicity during early neurodevelopmental periods and provide an age-dependent tauopathy similar to events that happened in the AD. The objective of this study was to determine the effects of ALA on the ERUPR signaling pathway in a human tau^R406W^ (htau) transgenic *Drosophila* model of tauopathy. We hypothesized that supplementation with ALA improves behavioral deficits and increases ERUPR efficiency via providing a better environment for neuronal survival. We showed that a lower dose of ALA modifies ERUPR and supported the reduction of behavioral deficits in 20 days old flies. As these beneficial effects were observed in 30 days old flies after treatment with a higher dose of ALA, we proposed that aging potentially interfere with ALA prophylactic effects.

## Methods

### Fly stocks and maintains

Fly strains used in this study and their sources were: *w*^*1118*^ (Bloomington *Drosophila* Stock Center (BDSC) at Indiana University #3605), *elav*-GAL4 (#8765), *ok*^*107*^-GAL4 (#854) and *gmr*-GAL4 (#8605). The *UAS-*htau^R406W^ transgenic stock (gifts from Feany MB, Harvard Medical School, Boston, USA) was used to induce tauopathy [24]. Heterozygous controls were obtained by crossing GAL4-drivers and UAS-effector to *w*^*1118*^. All the fly stocks were maintained on the standard diet media (1% agar, 6.5% cornmeal, 7.5% molasses, 1% yeast, and 0.3% propionic acid) under regular conditions (12h/12 h light/dark cycle, 25°C ± 1°C, and 50-60% relative humidity) in a vivarium where transferred to fresh maintenance diet after three days [25]. 25 males and 25 females flies were bulk propagated in 250 ml glass bottles having 25 ml of maintenance diet and the equal number of each gender was used for all experiments. Different flay strains (virgin females) were collected and allowed to mate and lay eggs according to the experimental design. Parents were put out from bottles after 24 h and eggs were allowed to hatch in a maintenance diet followed by larval pupation. F1 newly eclosed transgenic flies over 24 h, besides 10-, 20- and 30-days old flies have been collected for behavioral assessments as well as molecular and biochemical investigations.

### Study design and drug interventions

Several lines of evidence suggest that transcription of htau isoforms in transgenic flies can affect the developmental process of flies’ brain and sometimes observed neurodegeneration could be in part linked to developmental defects [26, 27]. In the present study, in order to minimize tau-induced neurodevelopmental defects, tau transcription was downregulated in both larval and early adult stages (days 1-10) via temperature-sensitive *UAS*/*GAL4* system [28]. In our model, transgenic flies were kept at 18 °C from the 1st instar larval stage up to the 10th day of an adult stage in order to reduce htau^R406W^ transcription in larvae and younger adults. In the next step, transgenic flies were kept in normal temperature (25 °C) from the 10th day to the 30th day of an adult stage in order to increase neurotoxicity of htau^R406W^ in older adults (adult-onset model). Alpha-lipoic acid (ALA, Santa Cruz Biotechnology, Dallas, USA) stock (0.25 mg/ml) was freshly prepared in a 1:8 solution of ethanol/PBS (pH 7.2) and added to adult flies diet in the final concentrations 0.001%, 0.005%, and 0.025% (w/w) during 10-20 or 20-30 days (figure 1). These concentrations of ALA were relatively in accordance with doses that are effective and extend lifespan in *Drosophila* [29]. Standard control diets contained 0.001%, 0.005%, and 0.025% (w/w) of 1:8 solution of ethanol/PBS as vehicle. Twenty-three male and twenty-three female flies of each experimental group in three replicates were used for gene and protein level analyses.

**Figure 1.**
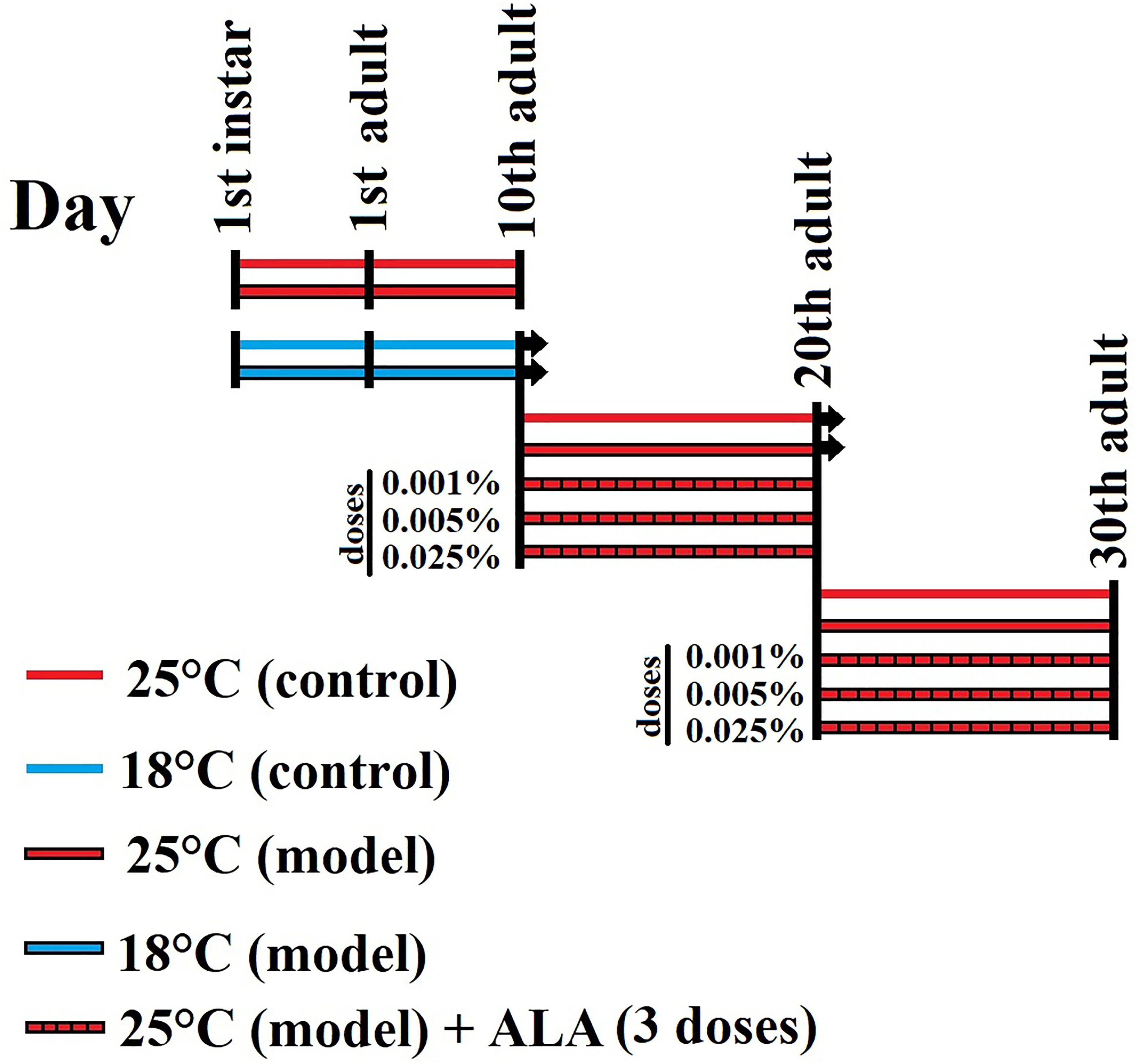
Experimental design, animal and treatment groups in this study.

### Experimental groups

In the model validation procedure, we had two experimental groups (all feed with normal diet) including a modified model group (growth in 18°C) and a traditional model group (growth in 25°C). Each group was consisting of corresponding control and model sub-groups (figure 1). After verification of model, we went to the drug investigating step with modified model flies and groups were consisting of: a) control group flies that feed with normal diet, transferred to 25°C after 10 days age, and undergo aging period (20 and 30 days adult flies), b) model group flies that feed with normal diet, transferred to 25°C after 10 days age, and undergo aging period, as well as c) treatment group flies that feed with normal diet contained of three doses of ALA, transferred to 25°C after 10 days age, and undergo aging period (figure 1).

### Flies eyes external morphology analysis

As the flies’ compound eyes structures are appropriate indicators of neurodegeneration, in order to examine whether eyes surface morphology is affected in the transgenic flies expressing htau^R406W^ in the compound eyes photoreceptor neurons by means of *gmr*-GAL4 driver micrographs of the external surface of the flies’ eyes were taken under a stereomicroscope. The htau^R406W^-*gmr* flies were CO2 anesthetized, immobilized on double-sided tape, and oriented laterally in a way that one eye exposed to the stereomicroscope objective lens. Images were captured under a white background and analyzed by ImageJ software. All phenotypic assessments of each tested genotype were analyzed in relation to the control genotype. The data were expressed as an average of 50 flies.

### Larvae olfactory conditioning for model confirmation

In both vertebrates and insects, olfactory information is first received by olfactory receptor neurons, which transmit signals into the CNS for future processing.

After activation of primary olfactory neurons by odorants, the first synapses in the complex neural networks called glomeruli are created. Subsequently, olfactory data are transported to higher brain centers, and finally perceived as odor and leads to behavioral response. As the corpora pedunculata (mushroom bodies) are essential brain structures for *Drosophila* olfactory memory, transcription of htau^R406W^ in mushroom bodies was achieved by making a genetic cross between *UAS*-htau^R406W^ and *ok*^*107*^-Gal4 (htau^R406W^-*ok*^*107*^). In order to confirm our model, the olfactory memory was assessed according to the standard protocol of larvae classical olfactory conditioning [30].

### Assay equipment’s and conditions

The late third instar larvae (96-102 h) old were separated by floating on 20% sucrose solution and washed in running tap water. We used a one-odor paradigm with an attractive odor (ethyl acetate) dissolved in mineral oil presented together with odorant vehicle control (mineral oil). The test and training plates filled with 0.01 M lithium chloride (to prevent drying and deserves uniform passage of the electric current) mixed with agarose (2%) to cover the whole bottom (1 cm thickness). Odorant and oil were loaded into 4.5-mm diameter Teflon containers with perforated lids and were used for maximum of 5 h post preparation. For equal distribution of odors during the test, the petri dishes were covered with parafilm. All experiments were done at room temperature (about 25°C) and parts of the experiments were done blind with respect to genotypes.

### Electroshock training and olfactory memory assay

In the naïve step, larvae (n=50) from each group were placed on the center of the petri dish (9 cm) covered by a layer of solidified agarose. 20 *μ*l of ethyl acetate added to one side and 20 *μ*l mineral oil added to another side of the petri dish and the larvae movements were monitored for 2 minutes. Larvae on each side of the petri dish were then counted, and a preference for ethyl acetate was calculated (preference of naïve (PN) = number flies in ethyl acetate side – number flies in oil side / total number of flies).

In the training step, larvae (n=50) from each group were placed on the center of the petri dish (9 cm) covered by a layer of solidified agarose. 20 *μ*l of ethyl acetate was dispensed on the inner surface of a 4.5 cm plate and was placed inverted on the plate where the larvae were present. Larvae are exposed to the ethyl acetate for 30 s and immediately they are exposed to the ethyl acetate and electrical shock (by passing pulses of 110 volts alternating-current (AC) between the electrodes) for another 30 s. This procedure was repeated three times (total time of 3 minutes). In the experiment step, two symmetrically opposite zones were distinguished by arcs of radius 2 cm. The arcs were drawn from the periphery of the petri dish (9 cm) containing a thin layer of solidified agarose. 20 *μ*l of ethyl acetate was poured in plastic micro cups and placed near the edge in one zone and 20 *μ*l of oil was poured in plastic micro cups and placed near the edge in another zone. Immediately after training, larvae were located at the center and the dish was covered. The time period of 2 minutes is given to the larvae to make the selection and larval quantification was made from video records. Larvae on each side of the petri dish were then counted, and a preference for ethyl acetate was calculated (preference of experiment (PE) = number flies in ethyl acetate side – number flies in the oil side / total number of flies). The performance index (PI) was calculated by subtracting the PN from the PE. The big values indicate higher olfactory memory, whereas the small values represent lower olfactory memory. The data were expressed as an average of three trials [30].

### Negative geotaxis assay for evaluation of ALA effects

The negative geotaxis assay in both htau^R406W^-*elav* and htau^R406W^-*ok*^*107*^ used for assessment of locomotor function as previously described [31]. Briefly, flies (30 adult males) were placed in a 25 cm length and 2 cm diameter, graduated empty plastic tube. After a short break (10 min), the flies were gradually tapped to the bottom of the tube and were allowed to moving upward to the top of the tube. The total number of flies (in each group) that after 1.5 min (every 10s) successfully climbed to 20 cm high of the tube were tracked by a blinded observer and analyzed. The data were expressed as an average of four trials per three independent replicates. Moreover, the raw video recordings were automatically analyzed via Ctrax software (http://ctrax.sourceforge.net/) and paths of movement, total velocity, and height climbed were reported [32].

### Quantitative reverse transcription PCR (qRT-PCR) analysis

In order to examine whether pan-neuronal transcription is affected in this model, the directed accumulation of htau^R406W^ was achieved from *UAS*-htau^R406W^transgene driven in the whole neurons with *elav*-GAL4 (htau^R406W^-*elav*). For qRT-PCR, forty-six flies heads (23 males and 23 females) were mechanically isolated (after snap freezing in liquid nitrogen) from each experimental groups and total RNA was extracted using an easy-BLUE total RNA extraction kit (iNtRON Biotechnology, South Korea) and resolved in 50μl DEPS water. Subsequently, cDNA was synthesized (from 5μg RNA) with a Maxime RT PreMix kit (iNtRON Biotechnology, South Korea) and real-time PCR was performed (from 1 μg of synthesized cDNA) using RealQ Plus Master Mix Green (Ampliqon, Denmark) according to the manufacturer protocol [33]. Amplification conditions were 95°C for 15 min (one cycle), 95°C for 20 sec (one cycle) and 65°C for 20 sec (45 cycles) followed by 72°C for 20 sec (one cycles). The used primer sequences were: Tau (F:5′-GGAAGACGTTCTCACTGATCTG-3′, R:5′-AGGAGTCTGGCTTCAGTCTCTC-3′), activating transcription factor 4 (ATF4) (F:5′-AGGCCATAGTACCCGCAAAC-3′, R:5′-CCGCCTGTTTGTAAGCATCG-3′), glucose-regulated protein 78kD/binding immunoglobulin protein (GRP87/Bip) (F:5′-GAATCAGTTGCACCAATCCC-3′, R:5′-AACTTGATGTCGTGTTGCACA-3′), X-Box Binding Protein 1 (XBP1) (F:5′-ACCAACCTTGGATCTGCCG-3′, R:5′-CGCCAAGCATGTCTTGTAGA-3′), ATF6 (F:5′-TGAGTTCCCCGATTTCTGGTT-3′, R:5′-GGAGACGAATTAGGCTCAGGT-3′), Ribosomal protein 49 (Rp49) (F:5′-TTCTACCAGCTTCAAGATGAC-3′, R:5′-GTGTATTCCGACCACGTTACA-3′). The average cycle threshold value (Ct) was calculated from three replicates per sample. Rp49 used as a reference gene and by 2^ΔΔ*Ct*^ method relative changes in gene expression were calculated [34].

### Western blot analysis

In order to examine whether the pan-neuronal protein level is affected in this model, the directed accumulation of htau^R406W^ was achieved through htau^R406W^-*elav* flies. Flies head were isolated as described in the qRT-PCR section and total protein was extracted using PRO-PREP kit (iNtRON Biotechnology, South Korea) according to the manufacturer protocol. Protein concentrations were estimated by the bicinchoninic acid (BCA) protein assay method (Sigma-Aldrich, Merck, USA). Proteins (50 μg) were resolved on 12.5 % SDS-PAGE gels and via electrophoretic transfer system (Bio-Rad, München, Germany) bonds moved to polyvinylidene fluoride (PVDF) membranes. After blocking of membranes with 1% (w/v) bovine serum albumin (BSA) they incubated (4°C overnight) separately with specific primary antibodies including rabbit anti-tau polyclonal antibody (1:500, Abcam, Catalogue no. ab64193), rabbit anti GRP87/Bip polyclonal antibody (1:600, StressMarq Biosciences, Catalogue no. SPC-180), mouse anti-ATF6 monoclonal antibody (1:750, Novus Biologicals, Catalogue no. NBP1-40256), rabbit anti-IRE1 (pSer724) polyclonal antibody (1:500, Novus Biologicals, Catalogue no. NB100-2323), rabbit anti-PERK polyclonal antibody (1:1000, Cambridge Research Biochemicals, Catalogue no. crb2005276), and mouse anti-α-Tubulin monoclonal antibody (1:1000, Cell signaling, Catalogue no. 2125). Next, membranes were carefully washed with PBS-T (PBS, 0.05% Tween-20), and incubated (4°C, 4 h) with horseradish peroxidase (HRP) conjugated antibodies against chicken, rabbit or mouse were obtained from Santa Cruz Biotechnology (Santa Cruz, CA, USA). The blots were exposed to DAB (3, 3′-Diaminobenzidine and H_2_O_2_) solutions for the detection of target antigens. Finally, after background subtraction and normalization of the density of each band to the density of tubulin, quantification of band intensities performed by ImageJ software (version 1.49) [35].

### Statistical analysis

GraphPad Prism software v.6 was used for all statistical analyses. Parametric statistics were performed for the groups that have not normal distribution (Shapiro–Wilk test) and homogeneity of variance (Bartlett’s test). To compare the effects of different temperatures and model validation data one-way analysis of variance (ANOVA) was applied followed by Bonferroni’s *post hoc* test (comparisons between groups larger than two). To compare the effects of different doses of ALA in different ages two-way ANOVA was applied followed by Bonferroni’s *post hoc* test. *P* values for levels of significance are represented as *< 0.05, **<0.01, ***< 0.001, and ****< 0.0001.

## Results

### Lower temperature significantly decreased htau^R406W^ neurotoxicity during brain development

As early brain development is very important for later health and nerve function we used the temperature-sensitive GAL4 system where minimum neurotoxicity of human tau^R406W^ observed. Our results showed that 10 days maintenance of flies in the low temperature (18°C) significantly decreases mRNA (Figure 2A) and protein (Figure 2B) level of htau^R406W^ (*P* < 0.001), and reduces the rough eye area (*P* < 0.05,) with no effect on the mean of eye size (Figure 2C) compared to the normal temperature (25°C) group. Moreover, no significant changes were observed in the preference index in larvae where growth in low temperature compared to the control group (Figure 2D). Furthermore, the htau significantly increases 10 days old flies’ locomotor impairment in both high temperatures (*P* < 0.001) and low temperatures (*P* < 0.05) compared to the control group (Figure 2E).

**Figure 2.**
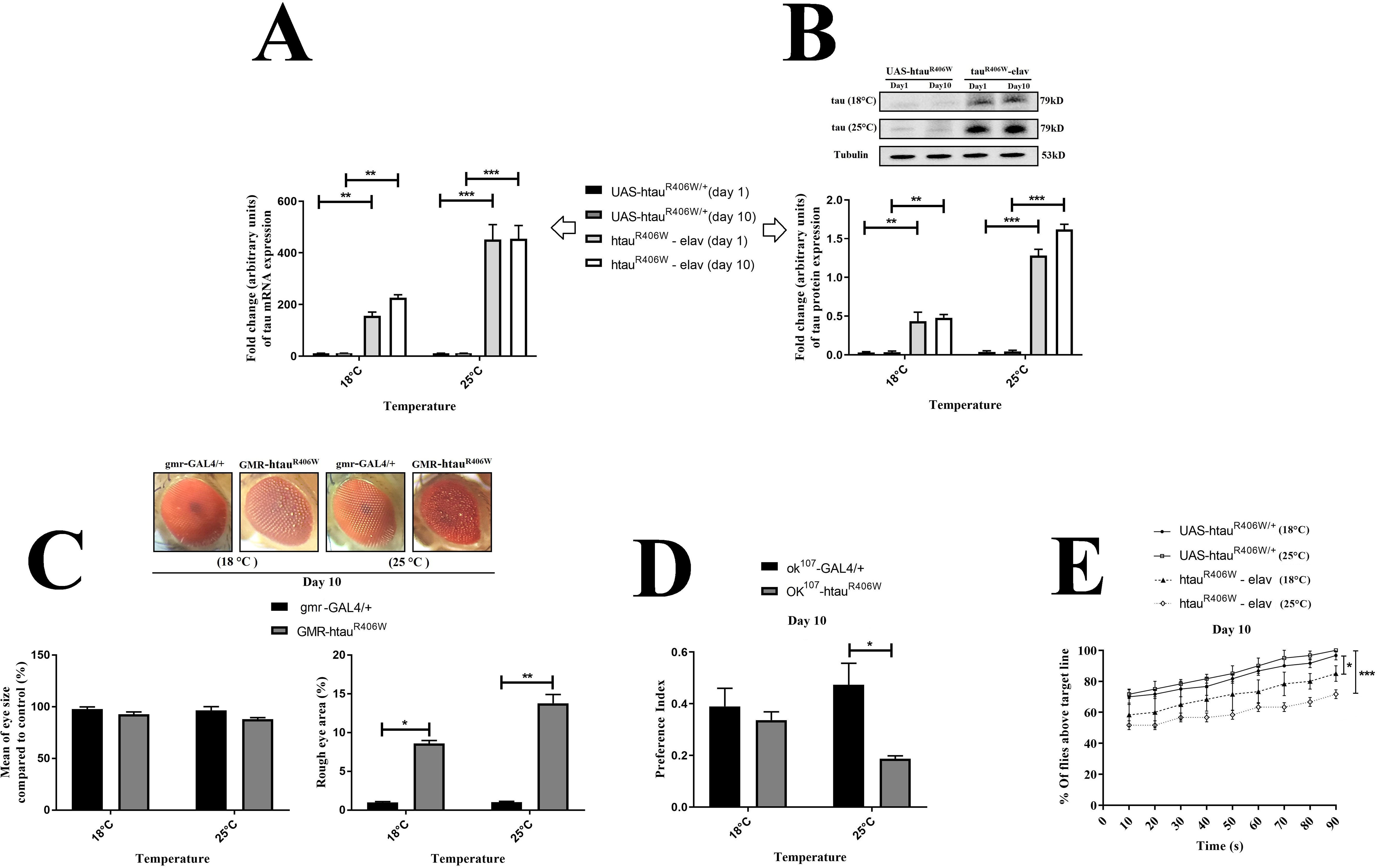
Molecular, morphological and behavioral evaluation of toxicity of human tau^R406W^ in the transgenic model of *Drosophila* under control of temperature-sensitive GAL4 system. Effects of low temperature (18 °C) in the mRNA (2A) and protein level of htau (2B). Effects of low temperature in eye external morphology (2C). Effects of low temperature in larvae olfactory memory (2D). Effects of low temperature in adult fly negative geotaxis (2E). Data analysis showed that 10 days maintenance of flies at a low temperature (18 °C) significantly decreased human tau^R406W^ toxicity via down-regulation of both mRNA and protein level of htau, improvement of eye morphology and reduction of behavioral deficits in larvae and adult flies in comparison with higher temperature (25°C) group. Data are presented as means ± SEM, analyzed using one-way ANOVA. *compared to relevant group, (^*^*p* < 0.05, ^**^*p* < 0.01, ^***^*p* < 0.001) with Bonferroni’s correction for multiple comparisons.

### Human tau^R406W^-induced ERUPR remains unchanged with aging

Although mRNA and protein levels of GRP87/Bip in the tauopathy group were significantly increased (*P* < 0.001) compared to the control group, aging did not significantly influence on mRNA and protein level of this key ER chaperone (Figure 3B and E). In the tauopathy group, expression of key ER stress sensors [XBP1 (Figure 4A), ATF4 (Figure 4B), and ATF6 (Figure 3C) in mRNA level and p-IRE1 (Figure 4C), p-PERK (Figure 4D), and ATF6 (Figure 3F) in protein level] were significantly increased (*P* < 0.001) compared to the control group, indicating activation of all three parallel signalling pathways of ERUPR (Figures 3 and 4). Changes in the protein and transcription of these sensors were not significant under the influence of senescence (Figures 3 and 4).

**Figure 3.**
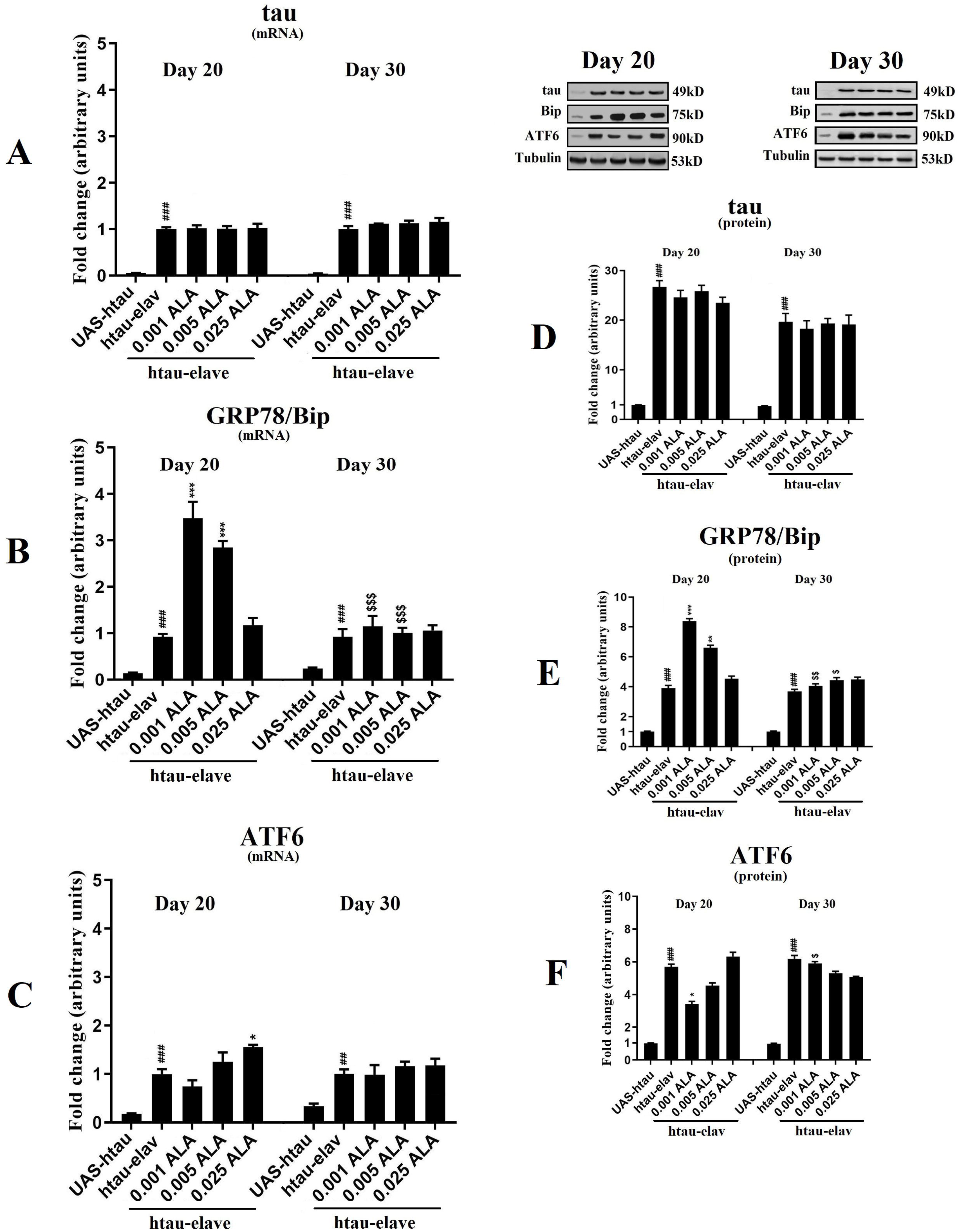
Evaluation of transcription and protein levels of htau (3A, 3D), GRP87/Bip (3B, 3E) and ATF6 (3C, 3F) in different ages and effects of ALA in these patterns in flies’ brains. Quantification of results shows that there are no significant changes in the htau transcription at all experimental groups. The transcription of both UPR markers (GRP87/Bip and ATF6) significantly increased in the tauopathy model in the comparison with control flies, and aging did not any significant effect on the transcription of these markers in both model and control flies. Gels are cropped and data are presented as means ± SEM, analyzed using two-way ANOVA. ^**#**^ compared to control group (*UAS-*htau) with same age, *compared to model group (htau-*elav*) with same age, ^**$**^ compared to same dose with different age (^*^, ^**#**^, ^**$**^*p* < 0.05, ^**^, ^**##**^, ^**$$**^*p* < 0.01, ^***^, ^**###**^, ^**$$$**^*p* < 0.001) with Bonferroni’s correction for multiple comparisons.

**Figure 4.**
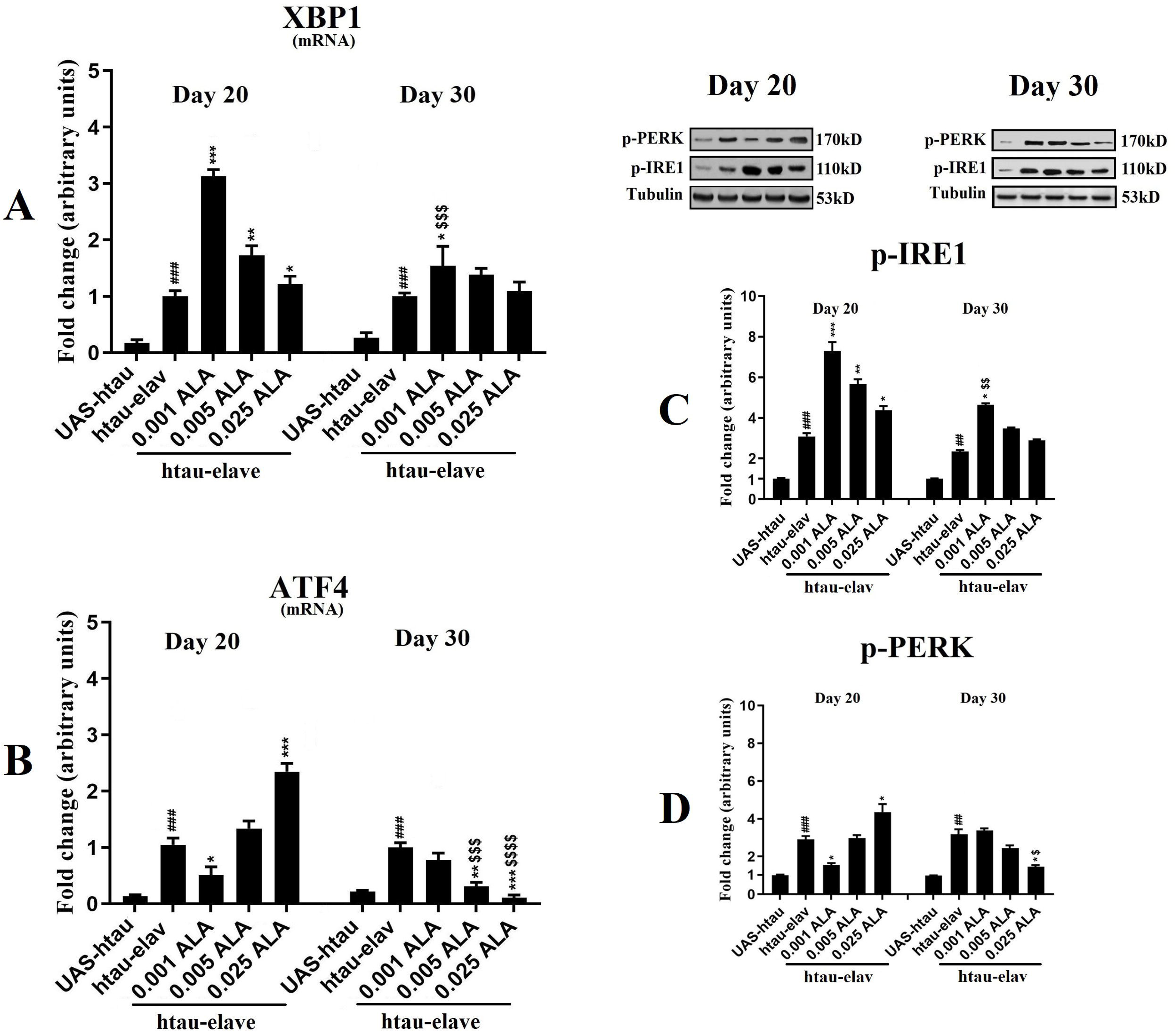
Evaluation of transcription of XBP1 (4A) and ATF4 (4B) as well as protein levels of p-IRE1 (4C) and p-PERK (4D) in different ages and effects of ALA in these patterns in flies’ brains. Quantification of results shows the dose-dependent effect of ALA on transcription of both XBP1 and ATF4 as well as expression of both p-PERK and p-IRE1 as key markers of ERUPR in the comparison with control flies, and aging has significant effect on these markers in both model and control flies. Gels are cropped and data are presented as means ± SEM, analyzed using two-way ANOVA. ^**#**^ compared to control group (*UAS-*htau) with same age, *compared to model group (htau-*elav*) with same age, ^**$**^ compared to same dose with different age (^*^, ^**#**^, ^**$**^*p* < 0.05, ^**^, ^**##**^, ^**$$**^*p* < 0.01, ^***^, ^**###**^, ^**$$$**^*p* < 0.001) with Bonferroni’s correction for multiple comparisons.

### Lower doses of ALA significantly increased GPR78/Bip only in the younger adults

The treatment with a low dose (0.001%) of ALA exhibited the most inductive effect on the protein level of major ER chaperone GRP87/Bip (*P* < 0.001) in younger (20 days) flies compared to the tauopathy group (Figure 3E). The results from treatment with middle dose (0.005%) of ALA also showed increases in GRP87/Bip gene and protein level (*P* < 0.001) in younger flies compared to the tauopathy group (Figure 3B and E). A higher dose (0.025%) of ALA treatment had no significant effect on gene and protein level of GRP87/Bip, in younger flies compared to the tauopathy group (Figure 3B and E). Remarkably, in older (30 days) flies, the expression of GRP87/Bip remained unchanged at both mRNA and protein levels after treatment with all three doses of ALA compared to the tauopathy group (Figure 3B and E). Noteworthy, the results showed that treatment with 0.001% and 0.005% doses of ALA significantly increased mRNA and protein levels of GRP87/Bip in younger flies (*P* < 0.001), compared to the older flies (Figure 3B and E).

### Lower doses of ALA significantly decreased ATF6 only in the younger adults

Although transcription of ATF6 was increased after high dose ALA treatment in younger flies, 0.001% and 0.005% doses of ALA did not significantly affect its transcription compared to the tauopathy group (Figure 3C). In the older flies, ATF6 transcription was not significantly amplified after treatment with all ALA doses compared to the tauopathy group (Figure 3C). The results showed that treatment with all doses of ALA insignificantly changed transcription of ATF6 in older flies, compared to the younger flies (Figure 3C). For a better understanding of ATF6 signaling, the effect of ALA treatment was investigated on the protein level of ATF6. The protein level of ATF6 was significantly reduced after low dose ALA treatment (*P* < 0.05) and insignificantly increased after high dose ALA treatment in younger flies compared to the tauopathy group (Figure 3F). On the other hand, the ATF6 protein level was insignificantly decreased after middle dose ALA treatment in younger flies compared to the tauopathy group (Figure 3F). In older flies, the ATF6 protein level was unchanged after treatment with all doses of ALA compared to the tauopathy group (Figure 3F). The results showed that treatment with a low dose of ALA significantly increased the protein level of ATF6 in younger flies, compared to the older flies (Figure 3F).

### ALA significantly increased activation of the IRE1 pathway in a dose-dependent manner

The ALA treatment exhibited a significant dose-dependent effect on the transcription of major IRE1 related transcription factor XBP1 in younger (20 days) flies compared to the tauopathy group (Figure 4A). A lower dose of ALA had a most significant effect (*P* < 0.001) and a higher dose of ALA treatment had a less significant effect (*P* < 0.05) on the transcription of XBP1, in younger flies compared to the tauopathy group (Figure 4A). Remarkably, in older (30 days) flies, the transcription of XBP1 significantly increased (*P* < 0.05) after treatment with 0.001 doses and insignificantly changed after treatment with 0.05 and 0.025 doses of ALA compared to the tauopathy group (Figure 4A). Noteworthy, the results showed that treatment with 0.001% of ALA significantly increased transcription of XBP1 in younger flies (*P* < 0.001), compared to the older flies (Figure 4A). According to the above-mentioned results, lover dose of ALA can significantly up-regulate XBP1 transcription in both younger and older flies compared to the tauopathy group (Figure 4A).

For a better understanding of IRE1 signaling, the effect of ALA treatment was investigated on the protein level of p-IRE1 (active form). The ALA treatment also exhibited a significant dose-dependent effect on the transcription of p-IRE1 protein in younger flies compared to the tauopathy group (Figure 4C). Lower had a most significant effect (*P* < 0.001) and a higher dose of ALA treatment had a less significant effect (*P* < 0.05) on the protein level of p-IRE1, in younger flies compared to the tauopathy group (Figure 4C). Remarkably, in older flies, the protein level of p-IRE1 significantly increased (*P* < 0.05) after treatment with 0.001 doses and insignificantly increased after treatment with 0.05 and 0.025 doses of ALA compared to the tauopathy group (Figure 4C). Noteworthy, the results showed that treatment with 0.001% of ALA significantly increased the protein level of p-IRE1 in younger flies (*P* < 0.01), compared to the older flies (Figure 4C).

### Age-dependent fluctuation in PERK pathway activation during ALA treatment

In order to examine the effects of ALA treatment on the PERK pathway activation, we evaluated major PERK related transcription factor ATF4 (in mRNA level) and p-PERK protein level in our experimental groups. As shown in figure 4B, ATF4 transcription was significantly reduced after low dose ALA treatment (*P* < 0.05) and significantly increased after high dose ALA treatment (*P* < 0.001) in younger flies compared to the tauopathy group. On the other hand, ATF4 transcription was not significantly changed after middle dose ALA treatment (0.005%) in younger flies compared to the tauopathy group (Figure 4B). Inversely, in older flies, the transcription of ATF4 significantly decreased after treatment with 0.025 (*P* < 0.001) and 0.005 (*P* < 0.01) doses and insignificantly changed after treatment with 0.001 dose of ALA compared to the tauopathy group (Figure 4B). Noteworthy, the results showed that treatment with 0.005% and 0.025% doses of ALA significantly decreased transcription of ATF4 in older flies (*P* < 0.01 and *P* < 0.001 respectively), compared to the younger flies (Figure 4B). For a better understanding of PERK signaling, the effect of ALA treatment was investigated on the protein level of p-PERK (active form). The protein level of p-PERK was significantly reduced after low dose ALA treatment (*P* < 0.05) and significantly increased after high dose ALA treatment (*P* < 0.01) in younger flies compared to the tauopathy group (Figure 4D). On the other hand, the p-PERK protein level was not significantly changed after middle dose ALA treatment in younger flies compared to the tauopathy group (Figure 4D). Inversely, in older flies, the p-PERK protein level only significantly decreased after treatment with 0.025% (*P* < 0.001) of ALA compared to the tauopathy group (Figure 4D). Noteworthy, the results showed that treatment with 0.001% of ALA significantly decreased the protein level of p-PERK in younger flies (*P* < 0.01), compared to the older flies (Figure 4D). On the other hand, treatment with 0.025% of ALA significantly decreased the protein level of p-PERK in older flies (*P* < 0.01), compared to the younger flies (Figure 4D).

### Effects of ALA on eyes morphology and locomotor dysfunction

We performed eye morphology analysis for ALA treatment groups of different ages (20 and 30 days), but there were no significant changes in eye-related factors (eye size and rough eye area) in the presence of ALA compared to the control group (data not showed). On the other hand, the tauopathy significantly increases (*P* < 0.001) locomotor impairment in both htau^R406W^-*elav* and htau^R406W^-*ok*^*107*^ flies in both 20 (Figures 5A1-4 and 6A1-4) and 30 (Figures 5B1-4 and 6B1-4) days old compared to the control group. Tauopathy model flies showed relatively same declines in the total velocity, paths of movement and height climbed indicated the same locomotor disability during the aging period (Figures 5 and 6). Low dose ALA notably increased the climbing ability of flies in the htau^R406W^-*elav* (*P* < 0.001) and htau^R406W^-*ok*^*107*^ (*P* < 0.01) youngest adults compared to the tauopathy group (Figures 5A1-4 and 6A1-4). Moreover, high dose ALA had not any significant effect on the climbing ability of youngest adult’s flies in both transgenic lines compared to the tauopathy group (Figures 5A1-4 and 6A1-4). On the other hand, treatment of both htau^R406W^-*elav* and htau^R406W^-*ok*^*107*^ flies with low dose ALA did not significantly affect behavioral parameters in the oldest flies compared to the tauopathy group (Figures 5B1-4 and 6B1-4). Remarkably, high dose ALA particularly increased the climbing ability of flies only in the htau^R406W^-*elav* (*P* < 0.01) oldest adults compared to the tauopathy group (Figures 5B1-4 and 6B1-4).

**Figure 5.**
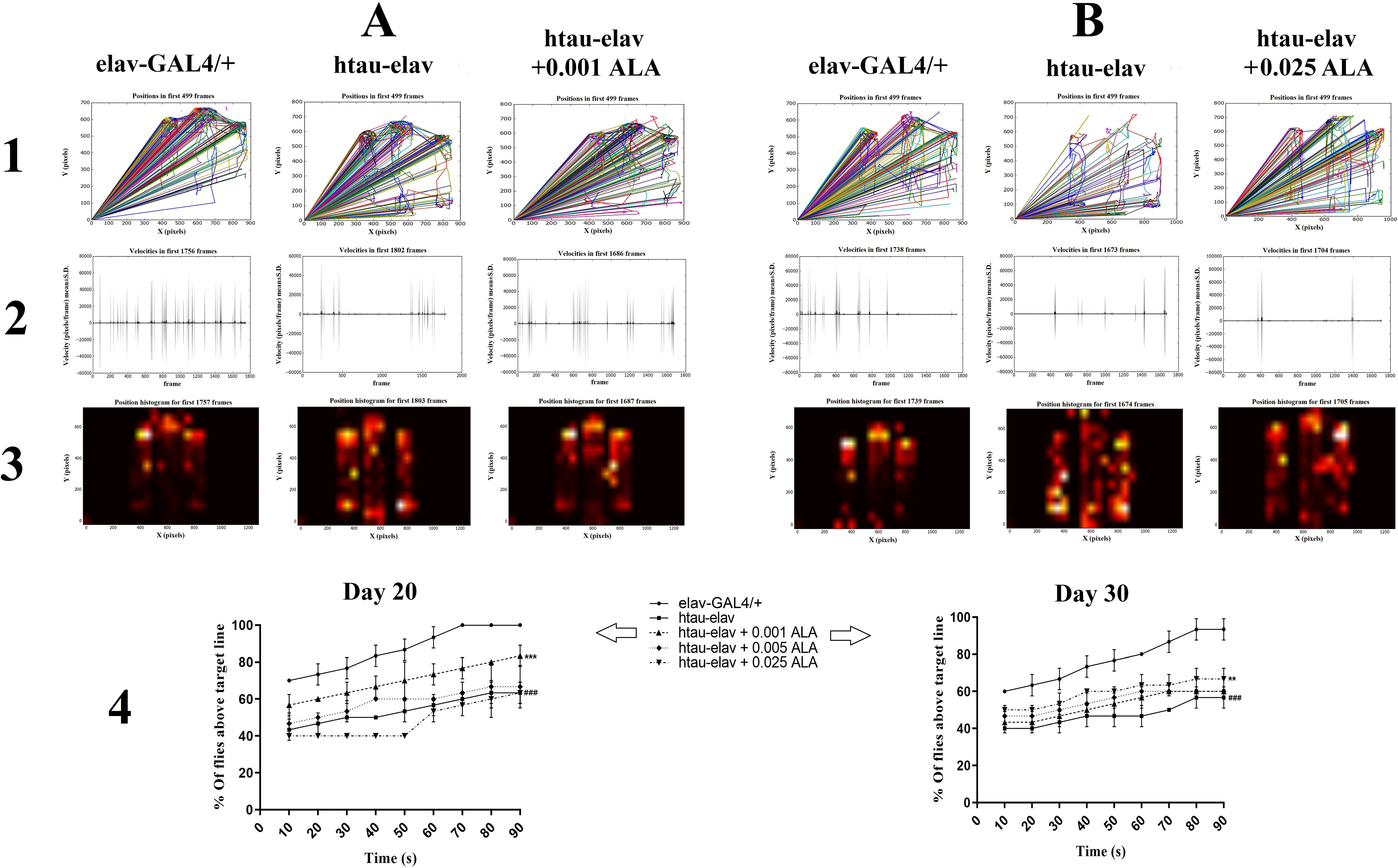
Evaluation of locomotor dysfunction of htau^R406W^-*elav* flies in different ages and behavioral effects of ALA. Negative geotaxis assays analyzed through Ctrax software and locomotor dysfunction in flies were evaluated. Ctrax outputs include: paths of movement (A1 and B1), total velocity (A2 and B2), and height climbed (A3 and B3). Negative geotaxis and ability to climbing also evaluated via analysis of movies (A4 and B4) as described in the method section. Quantification of results shows that locomotor dysfunction significantly increased in htau-*elav* flies in the comparison with control flies in both ages. Interestingly, only a lower dose of ALA (0.001) significantly decreased locomotor dysfunction on the younger adults in comparison with htau-*elav* flies (A4). On the contrary, only a higher dose of ALA (0.025) significantly decreased locomotor dysfunction on the older adults in the comparison with htau-*elav* flies (B4). Data are presented as means ± SEM, analyzed using two-way ANOVA. ^**#**^ compared to control (*elav-*GAL4/+) group, *compared to model (htau-*elav*) group (^**^ *p* < 0.01, ***, ^**###**^*p* < 0.001) with Bonferroni’s correction for multiple comparisons.

**Figure 6.**
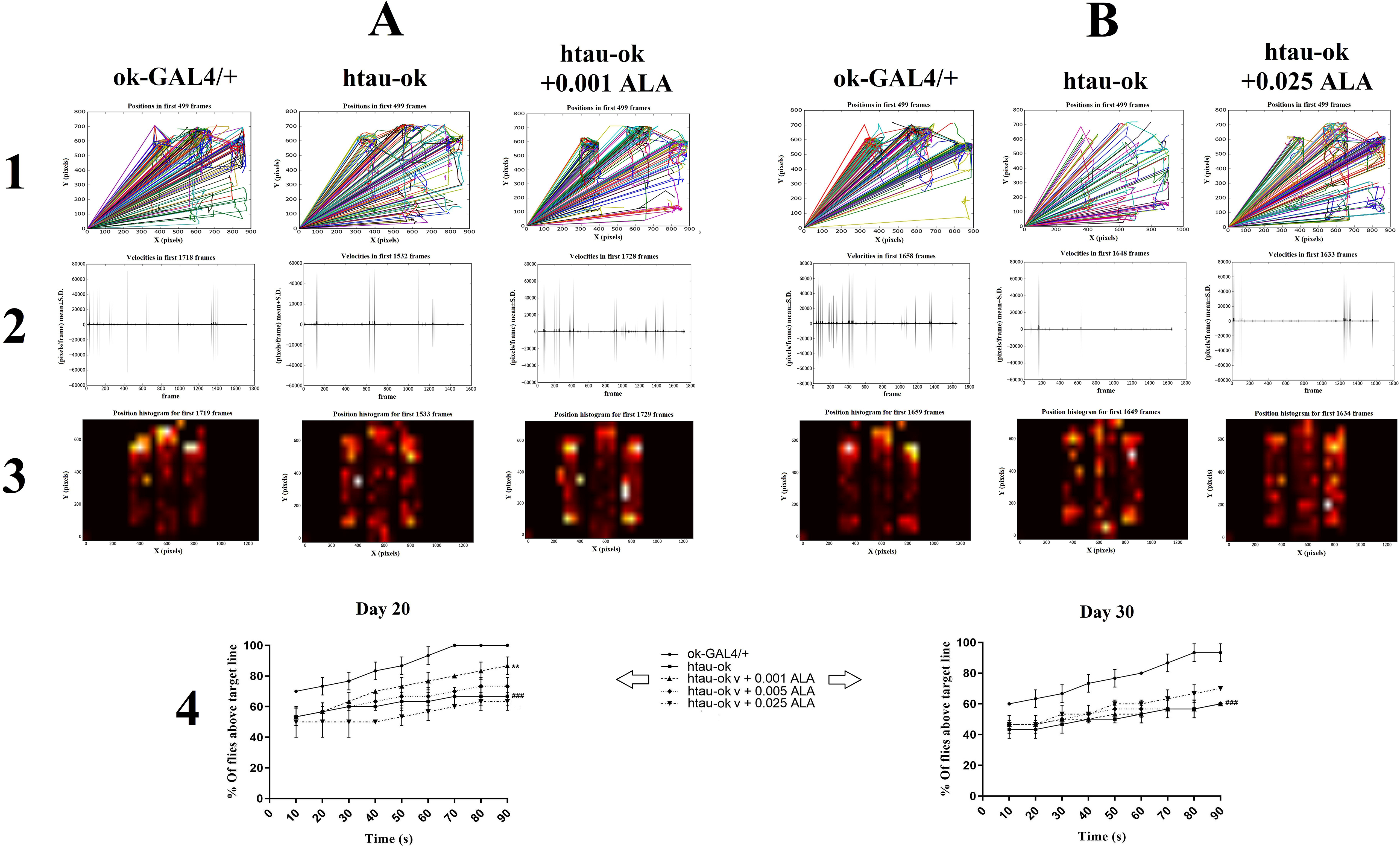
Evaluation of locomotor dysfunction of htau^R406W^-*ok*^*107*^ flies in different ages and behavioral effects of ALA. Negative geotaxis assays analyzed through Ctrax software and locomotor dysfunction in flies were evaluated. Ctrax outputs include: paths of movement (A1 and B1), total velocity (A2 and B2), and height climbed (A3 and B3). Negative geotaxis and ability to climbing also evaluated via analysis of movies (A4 and B4) as described in the method section. Quantification of results shows that locomotor dysfunction significantly increased in htau-*ok*^*107*^ flies in the comparison with control flies in both ages. Interestingly, only a lower dose of ALA (0.001) significantly decreased locomotor dysfunction on the younger adults in comparison with htau-*ok*^107^ flies (A4). On the other hand, ALA has no significant effect on locomotor dysfunction of the older adults in the comparison with htau-*ok*^*107*^ flies (B4). Data are presented as means ± SEM, analyzed using two-way ANOVA. ^**#**^compared to control (*ok*^*107*^-GAL4/+) group, *compared to model (htau-*ok*^*107*^) group (^**^ *p* < 0.01, ^**###**^*p* < 0.001) with Bonferroni’s correction for multiple comparisons.

## Discussion

Cellular processes involving transcription, translation, and folding of proteins, called proteostasis. Among them, protein folding is particularly important and is essential to its function. Proteostatic perturbations may establish by multiple factors, and frequently causing in the abnormal accumulation of misfolded proteins. Essential aspects of protein folding are cellular membrane dynamics; with could respond to the unexpected overload of secretory proteins inside the ER via mechanisms like the UPR. The cell’s ability to adjust the ERUPR to the toxic misfolded protein load finally dictates cell fate [36]. The main modifier of the cell proteostasis is the aging process, and the capability to conserve the equilibrium between folding and degradation of proteins, as well as the ability to respond to insults between these molecular pathways could disrupt during age-related tauopathies [37]. It has been demonstrated that the accumulation of tau stimulates the UPR by damaging ER-associated degradation (ERAD) proteins, including Hrd1 and valosin-containing protein [38].

In this study, we find that a low dose of ALA noticeably protected both htau^R406W^-*elav* and htau^R406W^-*ok*^*107*^ flies from neurotoxicity of htau in youngest adults via adaptation of ERUPR, which in turn enhances neuronal function and ameliorates locomotor function. The mechanism of action of low dose ALA in youngest adults involves the enhancement of GRP87/Bip, reduction of ATF6, downregulation of PERK-ATF4 pathway, and activation of the IRE1-XBP1 pathway. As a higher dose of ALA was barely able to improve locomotor function only in htau^R406W^-*elav* oldest adults but not in htau^R406W^-*ok*^*107*^ flies, we proposed that aging potentially affect ALA effective dose and mechanism of action.

The ER lumen harbors important molecular chaperones like GRP87/BiP that are recruited to misfolded nascent proteins for refolding in the correct structure. GRP87/BiP, a member of the heat-shock protein-70 (HSP70), is a sensor master to initiate ERUPR via the three main signaling pathways [39]. Under normal conditions, GRP87/BiP is localized to the ER lumen, and ATF-6, PERK, and IRE1 remain in a state of inactivity due to binding of GRP87/BiP [40]. Upon ER stress, the formation of misfolded proteins inhibits the interaction between GRP87/BiP and mentioned ER stress sensor proteins, thereby initiating ERUPR signaling pathway [41]. In our study, we showed that a lower dose of ALA significantly increased GRP87/BiP as a negative regulator of ERUPR in the youngest adults. On the other hand, all three doses of ALA were unable to change the level of GRP87/BiP in the oldest adults. As, up-regulation of GRP87/BiP, is important for proper folding of partly folded and newly synthesized proteins to maintain protein homeostasis, for the first time we demonstrated that aging processes diminish neuroprotective effects of ALA in the human tau^R406W^-induced ERUPR.

Moreover, IRE1 is the most evolutionarily conserved sensor, which undergoes dimerization and trans-auto-phosphorylation upon ER stress, and cleaves off a 26 nucleotides intron from mRNA encoding XBP1. The XBP1 upregulates genes involved in ER and Golgi biogenesis and cells lacking XBP1 are more sensitive to hypoxia-induced apoptosis [39]. In addition, activation of important UPR sensor IRE1 also stimulates stress-related kinases, including c-Jun N-terminal kinase (JNK protein) through bind to tumor necrosis factor receptor-associated factor 2 (TRAF2) [40]. It has been reported that the active form of XBP1 (XBP1s) up-regulates numeral genes that regulate protein quality control or misfolding [42]. In some neurodegenerative disorders, stimulation of UPR activation, as well as temporary and modest induction of the IRE1/XBP1 pathway, has neuroprotective effects [43, 44]. In our study, we showed that ALA significantly increased IRE1-XBP1 activation in the youngest adults and ameliorates behavioral deficits in a dose-dependent manner. Here, for the first time, we demonstrated that aging processes decline the effects of ALA on IRE1-XBP1 activation in this model of tauopathy.

PERK after activation gets trans-auto-phosphorylation and homodimerized, thus stimulating the phosphorylation of cytoplasmic eIF2α at serine residues. Phosphorylated eIF2α, induced the production of specific mRNAs bearing an internal ribosome entry site (IRES), like the ATF4 [40]. ATF4 controls ER redox homeostasis, protein folding, amino acid biosynthesis, cholesterol metabolism, and autophagy, but also the fate of a cell through the induction of the CCAAT enhancer-binding protein (C/EBP) homologous protein (CHOP) and members of the BCL□2 (B□cell lymphoma 2) protein family. Reduction in the phospho-eIF2α levels through induction of protein phosphatase GADD34 (the growth arrest DNA damage□inducible protein 34) and reversion of the inhibition of cellular global translation could also be mediated by CHOP [45]. ATF4 translation is a target of UPR and the integrated stress response through phosphorylation or eIF2α. Therefore, several kinases targeting eIF2α could be responsible for ATF4 levels modulations. The results of this study showed a strong correlation between p-PERK and ATF4 levels, suggesting that PERK is probably the main regulator of ATF4 mRNA levels in tauopathies. In our study, we showed that a lower dose of ALA significantly decreased but a higher dose of ALA significantly increased PERK-ATF4 activation in the youngest adults. On the other hand, only higher doses of ALA were able to significantly deactivate PERK-ATF4 in the oldest adults. Here, for the first time, we demonstrated that aging processes reversed the effects of ALA on PERK-ATF4 deactivation in this model of tauopathy.

ATF6, a type II ER-transmembrane protein with a bZIP domain (basic leucine zipper DNA-binding domain), besides two other UPR transducers PERK and IRE1, which are type I transmembrane proteins, serves as key sensors of the UPR signaling in the ER. During the oxidative stress, the luminal domain of ATF6 loses its association with GRP87/Bip and causes translocation of ATF6 from the ER to the Golgi [39, 46]. In our study, we showed that low dose ALA exerts their protective effects in youngest adults via decreasing ATF6 protein level.

Several experimental studies have indicated the neuroprotective effects of ALA in neurodegenerative diseases with oxidative stress, such as AD and Parkinson’s disease (PD). For example, Quinn et al. reported that chronic supplementation with ALA (0.1 % w/w of diet) enhanced memory and learning in the Morris water maze and reduced hippocampal memory deficits in transgenic aged Tg2576 mice as a model of cerebral amyloidosis associated with AD [47]. Another study by Fava et al. showed that ALA (600□mg/day) could be an effective therapeutic agent in slowing cognitive reduction in AD patients [48]. It has been reported that chronic (10 months) ALA administration had smaller effects on Y-maze performance on the young and aged mice and did not decrease end-point amyloid-β load, but significantly reduced in markers of oxidative stress [49]. These results were in accordance with our finding and suggest that despite the clear antioxidant property of ALA, chronic ALA therapy, at levels within tolerable nutritional guidelines were reduce oxidative modifications, and have limited benefits in aging and AβPP-transgenic mice [49]. Another study evaluated the effects of ALA in senescence-accelerated mouse prone 8 (SAMP8) model and indicated that ALA could improve memory and reverses oxidative stress in extremely old SAMP8 mice. The results from the Barnes maze test indicated that ALA-treated mice made fewer errors and spent more time by the target than the control groups, but decreased lifespan [50]. It has demonstrated that ALA (50 and 100 mg/kg) improves motor impairment related to the 6-hydroxydopamine-induced model of PD [20].

It has been also shown that ALA reverses the age-associated loss of neurotransmitters and their receptors, which can underlie its effects on memory deficits related to aging and neurodegeneration [51]. Li et al. also evaluated the effects of ALA in lipopolysaccharide (LPS)-induced inflammatory PD model, and reported that ALA treatment (100 mg/kg/d) improved motor dysfunction [52]. In the other study, ALA supplementation (0.23% w/v in drinking water for 4 weeks) led to important changes in synaptic function and was more effective in reversing the impaired synaptic plasticity in the old mice, than young mice [53]. In the more relative research ALA (200 μM) significantly attenuated glutamate-induced ER-stress markers; namely, GRP87, ATF6, PERK, eukaryotic translation initiation factor 2 alpha (eIF2α), RE1, CHOP, and caspase-12 in C6 Glioma cells [54]. It also has also reported that ALA induced transcription of ER stress-related genes, such as GRP87, CHOP, and XPB1s in A549 cell lines [18]. It has been demonstrated that in Drosophila, PERK and IRE1/XBP1 signaling (canonical UPR pathways), were induced upon ER stress during aging. In addition, transcriptions of many UPR-related genes were significantly elevated in aged brains, suggesting the activation of ERUPR in aged flies’ brains. Conversely, ERAD activity was reduced after activation of the ERUPR in aged tissues, signifying that the ERAD machinery may be compromised at a post-transcriptional level [55]. Anyway, we should mention that our transgenic model more resembles primary age-related tauopathy (PART) versus Alzheimer’s neuropathology, which is frequently observed in the brains of aged individuals, where tau-positive NFTs is directly associated with cognitive deficits even in the lack of Aβ plaques [56, 57].

### Limitations of the study

We observed that the effects of ALA on UPR markers are very dissimilar with dose independent-manner. ERUPR is a multipart mechanism that several known or unknown molecules take part in this response. These molecules could potentially affect each other on the feedback-positive or - negative molecular pathways. For example, dimerization of PERK with GRP87/Bip happened in resting cells and oligomerization occurred in cells under ER-stress [58]. Thus, ALA could have either directly or indirectly affected the EUPR mediators. We propose that suppression of tau-induced PERK phosphorylation and ATF4 expression in younger flies could mediate with downstream signaling of GRP87/Bip which up-regulated with ALA. On the other hand, suppression of tau-induced ATF6 expression could either mediate with GRP87/Bip or IRE1 where ALA stimulated IRE1 phosphorylation and induced XBP1 expression in younger flies. In this regard, here we did not investigate if inactivating GRP87/Bip or IRE1, that supposed to are the main mediators for ALA effects, with either antagonizing or gene silencing strategies could result in a significant reduction in ALA effects or not. Finally, it would have been a good idea to assess the effect of ALA on transgenic flies without GRP87/Bip or IRE1. This approach could provide new funding regarding the exact molecular mechanism of ALA action in this model. We also propose that the evaluation of lower doses of ALA in this model could potentially shed light on the neurotoxicity and effective dose of ALA.

## Conclusions

Controversies in the literature over the effects of ALA in the ER-stress, dose-independent effects of ALA, and lack of information about the role of aging in ER function during tauopathy make it difficult to explain the exact molecular mechanism of ALA effects in our study. Altogether, in this study, we highlighted the protective properties of ALA in the tauopathies in light of its effects on ERUPR and its implication in the aged brain. Using this transgenic model of tauopathy for avoiding of developmental effects of toxic tau, flies displayed behavioral alterations and activation of the three arms of the UPR. ALA administration improved behavioral deficits and modulated UPR activation induced by tau toxicity more efficiently in young adults, which suggests an age-dependent decline in the response. These observations, for the first time, raise questions about the role of ALA in the tauopathy and provide preliminary evidence that senescence-related cellular processes may affect the benefits of ALA. Additionally, our data reflecting the human tau^R406W^-induced ERUPR would be a good model for future researches in this kind. The implications of the association of ALA doses with its adverse effects and their mechanism of action also remain to be clarified, although in this case a principal role for aging processes has been proposed. Determining pathobiology of tau neurotoxicity in *Drosophila* aged brains, with recognition of key molecular mediators, will provide a firm basis for a more thoughtful understanding of age-related tauopathy.

## Declarations

### Ethics approval and consent to participate

N/A.

### Consent for publication

N/A.

### Availability of data and material

Data related to this study are either presented in the results section. Raw data can be obtained from the corresponding author on reasonable request.

### Authors’ contributions

EZ-G and NS conceived and designed the experiments, interpreted the data and drafted the manuscript. EZ-G performed all the experiments as their PhD thesis. KP and GV offered valuable suggestion and helped in drafting of manuscript. All authors read and approved the final manuscript.

## Acknowledgements

The authors are grateful to all respected research staffs in the Department of Biology, University of Zabol (especially Dr. Mohammad Haddadi), for their help with the study.

## Abbreviations

AD: Alzheimer’s disease
ALA: Alpha-lipoic acid
APP: Amyloid precursor protein
ATF4: Activating transcription factor 4
ATF6: Activating transcription factor 6
GRP87/Bip: Glucose-regulated protein 78kD/binding immunoglobulin protein
IRE1: Inositol regulating enzyme 1
MAPT: Microtubule-associated protein tau
NFTs: Neurofibrillary tangles
PERK: Protein kinase RNA-like ER kinase
Rp49: Ribosomal protein 49
UPR: Unfolded protein response (
XBP1: X-Box Binding Protein 1

